# Free energies of the Gln tautomerization and rotation mechanism of dark-state recovery in blue light-using flavin proteins

**DOI:** 10.1101/2023.08.11.551373

**Authors:** Alberto Pérez de Alba Ortíz, Carme Rovira, Bernd Ensing

## Abstract

Blue light-using flavin (BLUF) proteins are light-sensors that regulate responsive movement, gene expression and enzyme activity in diverse organisms. Their signaling times range from seconds to minutes, indicating a uniquely flexible dark-state recovery mechanism. Unlike other light-sensors, the flavin chromophore is non-covalently bonded to the protein. Hence, the switching occurs via a change in the protein-flavin hydrogen-bond network, involving conserved residues transferring protons, tautomerizing, rotating, and approaching or leaving the chromophore pocket; triggering secondary structure displacements. The specific deactivation steps and residue roles have remained controversial. The detailed process is difficult to probe experimentally, and although simulations can track it, the computational effort is daunting. We combine forefront techniques to simulate, for the first time, explicit dynamics of the deactivation. A hybrid quantum mechanics/molecular mechanics scheme focuses the computational resolution in the flavin’s vicinity, while our path-based methods sample the mechanism of dark-state recovery with high efficiency. Our protocol delivers free-energy profiles for the deactivation of two BLUF proteins, BlrB and AppA; corroborating a proposed mechanism based on the rotation and tautomerization of a conserved Gln. We find that the conformation of a Trp and a Met near the flavin is crucial to modulate the rate-determining barrier, which differs significantly between the BlrB and AppA proteins. Our work evidences how specific variations of the deactivation mechanism control vast differences in signaling times.

## 1 Introduction

The importance of being able to respond to external stimuli has driven the evolution of light-sensing systems in all kingdoms of life. Different kinds of photoreceptor proteins have been discovered across species; each binding a particular chromophore, i.e. a molecule that absorbs photons of a specific wavelength [71]. The photoreceptor protein converts the energy absorbed by the chromophore into a signal, which is propagated via chemical and conformational changes; ultimately triggering a biological response. Blue light-using flavin (BLUF) photoreceptors, discovered in 2002 [35, 50, 22], have a flavin-based chromophore— flavin adenine dinucleotide (FAD)—and are found in many bacteria and in some algae. Among other functions, BLUF proteins regulate phototaxis, as well as photosynthetic gene expression [14]. Unlike the chromophores of other photoreceptors, such as rhodopsin, which undergo cis-trans isomerization upon illumination [71], the flavin does not present significant conformational changes under blue-light. In contrast, BLUF activation is characterized by a change in the hydrogen-bond network between the FAD and a few conserved protein residues, which is caused by photo-induced proton-coupled electron transfers (PCETs). This hydrogen-bond rearrangement is completed within *<* 1 ns after photoactivation, and followed by a conformational change that propagates the signal to the outside of the protein in *µ*s to ms [54, 14]. The signaling time, which is determined by the rate of the thermal return to the dark state, can vary from a couple of seconds to several minutes across different BLUF domains, e.g. ∼ 2 s in BlrB [76], ∼ 26 s in PixD and ∼ 25 min in AppA [16, 17]. Several signaling mechanisms have been proposed, however the control of these signaling times in the different members of the BLUF family has remained thus far unclear. A deeper understanding about the mechanistic origin of the widely-varying recovery times of BLUF proteins could enable a precise control of their photoactivation in optogenetic tools [7] and biosensors [67].

Determination of the different *light*- and *dark*-state hydrogen-bonding patterns in BLUF proteins has demanded arduous experimental and computational efforts [14, 49]. Early spectroscopic analyses indicated stronger hydrogen-bonding to the C_4_ O carbonyl of the FAD in the signaling state [48]. X-ray crystallography identified Tyr, Gln and Asn as the conserved residues interacting with the C_4_ O carbonyl of the FAD. Mutagenesis studies [46, 45] showed that only Tyr and Gln are essential to form the signaling state; confirming the key hydrogen-bond network FAD-Gln-Tyr. However, the exact light- and dark-state hydrogen-bonding motifs remained elusive and debated for more than a decade, with models proposing the conserved Gln undergoing either a rotation [1], an amide-imide tautomerization [61], or both [10, 43, 69] to explain the stronger C_4_ O hydrogen-bonding upon activation [14]. Very recent efforts combining Fourier-transform infrared (FTIR) spectroscopy and quantum chemical calculations revealed that the activation is given by both rotation and tautomerization of the conserved Gln [11, 36]. In Fig. 1 we show the structures assigned to the light and dark states. In the light state, the Gln is in the ZZ-imide form, with its O-H group hydrogen-bonded to the C_4_ O carbonyl of the FAD. In the dark state, the Gln is rotated ∼ 180° with respect to the light state and in the amide form, with its oxygen hydrogen-bonded to the Tyr. In both states, the nitrogen of the Gln forms a hydrogen-bond to the N_5_ of the FAD. Further computational studies have validated the stability of these hydrogen-bond networks [20], as well as demonstrated the feasibility of the PCETs that facilitate the Gln tautomerization upon excitation [68, 21, 63, 74, 19]. Femtosecond time-resolved absorption studies also support the PCET, but only with Gln tautomerization in PixD[15], whereas recent polarizable QM/MM calculations of diverse spectra support the PCET, as well as both Gln tautomerization and rotation in AppA[29, 27].

**Figure 1:**
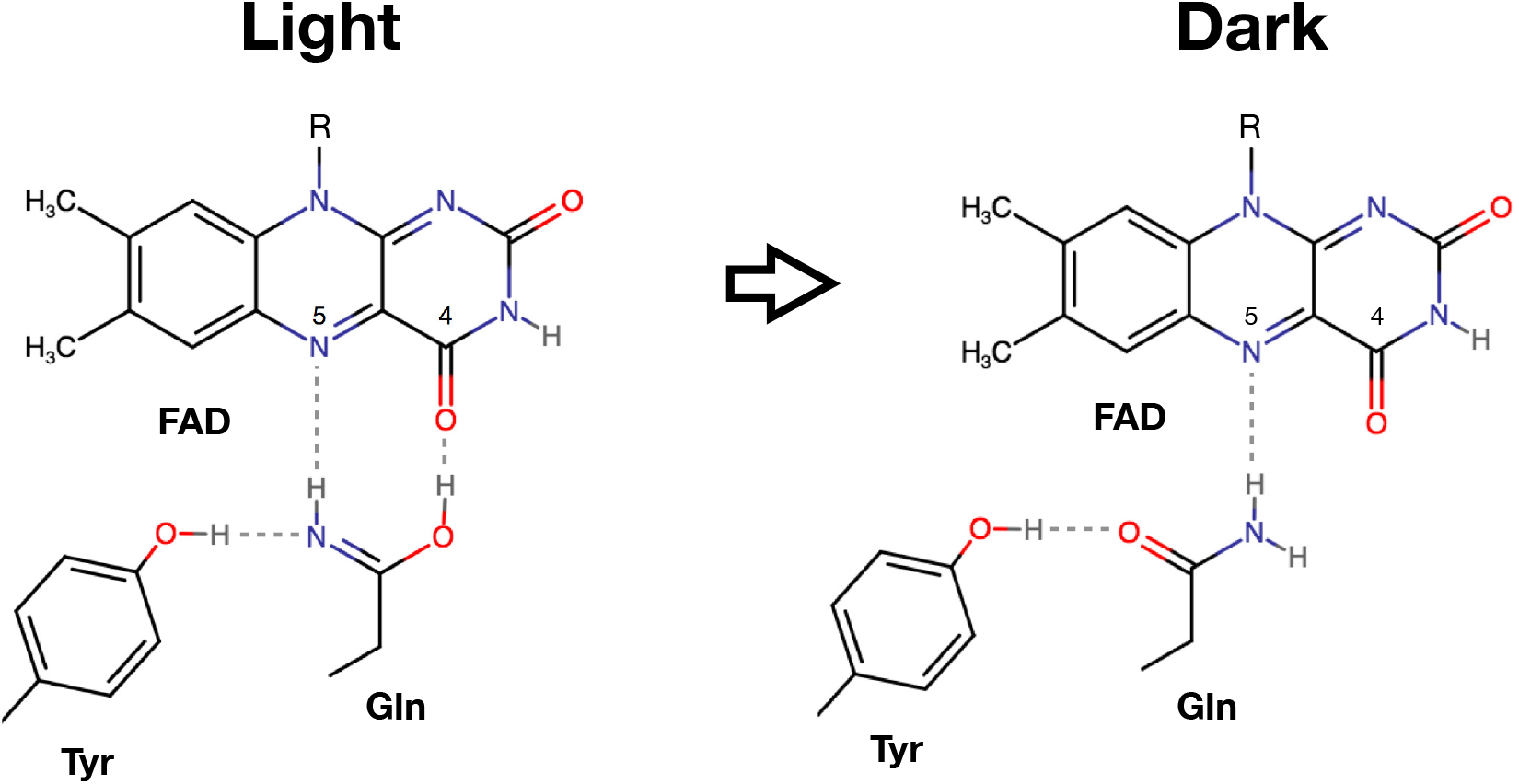
Light- and dark-state structures of the Tyr-Gln-FAD hydrogen-bond network in BLUF proteins. The thermal return to the dark state is achieved via Gln rotation and tautomerization from ZZ-imide to amide form.

After the hydrogen-bond rearrangement within the active site, the light-induced signal is propagated via conformational changes across the BLUF protein. Two highly conserved residues located in the fifth *β* strand, Met and Trp, have been identified as key components in the signal transduction mechanism (see Fig. 2). Ultimately, the signal is transmitted to the C-terminal region, which has been shown to control slow dynamics of BLUF domains [66]. In the PixD protein, replacing the conserved Met with Ala does not hinder the FAD-Gln-Tyr hydrogen-bonding change, but prevents further conformational changes toward the C-terminal helices, as well as the usual regulation of phototaxis in *Synechocystis* cyanobacteria [51]. Similarly, mutating Trp to Ala in the AppA protein also abolishes conformational changes in the *β* strand [52], as well as the light-responsive photosynthetic gene expression in *Rhodobacter sphaeroides* [53]. Interestingly, replacing Trp for Ala in PixD does not have significant effects on the conformational switch, or in the downstream phototaxis [51]; indicating that the roles of Met and Trp change depending on the BLUF domain. Directed changes in the H-bond networks of AppA can make its photocycle similar to that of PixD [30]. Recent Förster resonance energy transfer (FRET) measurements indicate that, in AppA, the Trp is rigidly hydrogen-bonded near the FAD in the light state, and flexible and pointing away from the FAD in the dark state [41]. Based on X-ray crystallography, such Trp_in_/Met_out_ [1] (light) and Trp_out_/Met_in_ [40] (dark) conformations had already been proposed for the AppA protein. Computational studies have also been dedicated to elucidate the Trp/Met switching mechanism in AppA [23, 21]. In contrast, the Trp/Met switch does not occur in other BLUF domains, such as BlrB, where Trp faces outward in both states [36]. Recent studies using mutation and femtosecond spectroscopy have proposed unified elementary steps for AppA and another two BLUF domains, *Oa*PAC and *Sy*PixD [75], but not with BlrB. These mechanistic variations between BLUF domains might hold the key to understanding their wide span of dark-state recovery times.

**Figure 2:**
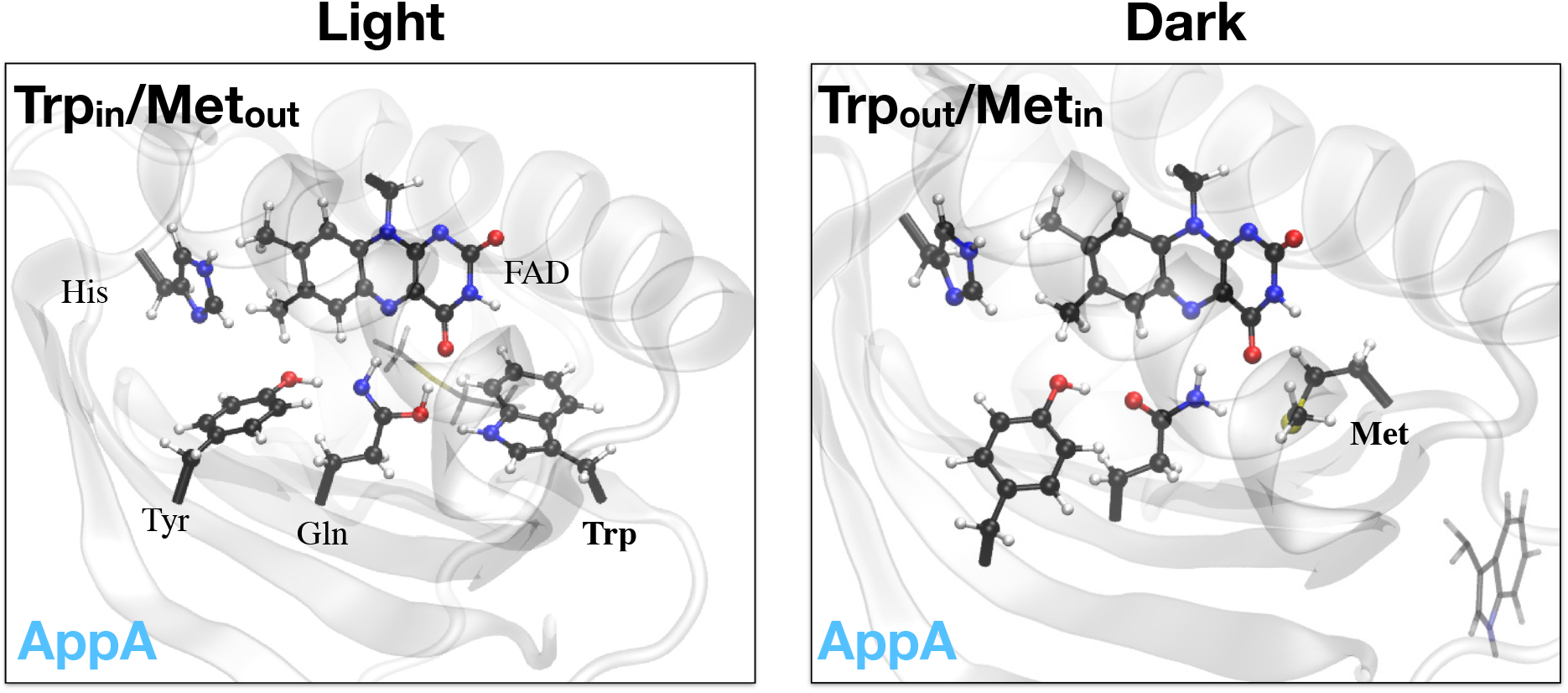
AppA BLUF protein in the light state, with the Gln in imide-ZZ form and the Trp_in_/Met_out_ conformation; and in the dark state, with the Gln rotated in amide form and the Trp_out_/Met_in_ conformation. Structures are taken from MM simulations of the 1YRX [1] and 2IYG [40] PDB structures, respectively. In the BlrB protein, the transition to the dark state also involves Gln rotation and tautomerization, while remaining in the Trp_out_/Met_in_ conformation [36].

The experimentally and computationally validated structures enable the investigation of light-to-dark pathways, whose mechanistic details can explain the differences in recovery rates across BLUF domains. Hybrid quantum mechanics/molecular mechanics (QM/MM) calculations [73, 12, 64] are especially well-suited to treat the deactivation process in BLUF photoreceptors [56, 62]. The vicinity of the FAD, where the proton transfers occur, is described at the more accurate—but costly—QM level, i.e. with density functional theory (DFT); while the surrounding protein and solvent environment is simulated at the more affordable MM level, i.e. with a force field. Most of the previous computational studies about specific steps of the BLUF photoactivation involve QM/MM calculations [61, 10, 69, 43, 36, 23, 21, 74, 44], and some studies have focused on the effect of the selected QM/MM parameters [70, 55, 28]. Shortly after the first FTIR spectroscopy and QM simulation confirmation of the Gln tautomerization and rotation [11], Khrenova and coworkers proposed a mechanism for the dark-state recovery in the BlrB BLUF domain [42]. The mechanism involves a nearby conserved protonated His and is comprised of the following steps (see Fig. 3): 1) concerted proton transfers from the nearby conserved protonated His to the Tyr, and from the Tyr to the nitrogen of the Gln; 2) ∼ 180° rotation of the O-H hydroxyl group of the Gln, which breaks the hydrogen bond with the C_4_ O carbonyl of the FAD; 3) ∼ 180° rotation of the Gln; and 4) concerted proton transfer from the oxygen of the Gln to the Tyr, and from the Tyr to the nearby His, which complete the imide-to-amide tautomerization of the Gln. The Met on the fifth *β* strand remains within the active site throughout the transition, as suggested by experiments [36]. Using hybrid QM/MM calculations, Khrenova and colleagues obtained potential-energy profiles for the recovery of the BlrB protein and its Tyr9^3F^ mutant, in which the conserved Tyr is mono-fluorinated. The BlrB potential-energy profile—with barriers of 13.2 and 14.6 kcal/mol for steps 2) and 3), respectively—implicated a recovery time of 0.4 s, close to the 2 s reported experimentally [76]. Fluorination of the conserved Tyr, which lowers its p*K*_a_, increased the potential-energy barriers by ∼ 4 kcal/mol, corresponding to a theoretical recovery time of 4.8 s. The lengthened recovery time predicted for the BlrB Tyr9^3F^ mutant agreed qualitatively with reported experiments for the PixD BLUF protein [54]. More recent experiments suggest that the trend might be opposite, with a recovery time of 26 s for the PixD wild type and 4.5 s for its Tyr9^3F^ mutant [17]. However, such a difference in the rate would amount to a change of ∼ 1 kcal/mol in the calculated barrier, which lies arguably beyond the accuracy limit of this sort of QM/MM energy calculations [32]. Recently, the rotated and tautomerized Gln has also been observed in QM/MM free-energy perturbation calculations[65] and in polarizable QM/MM calculations of various spectra [27].

**Figure 3:**
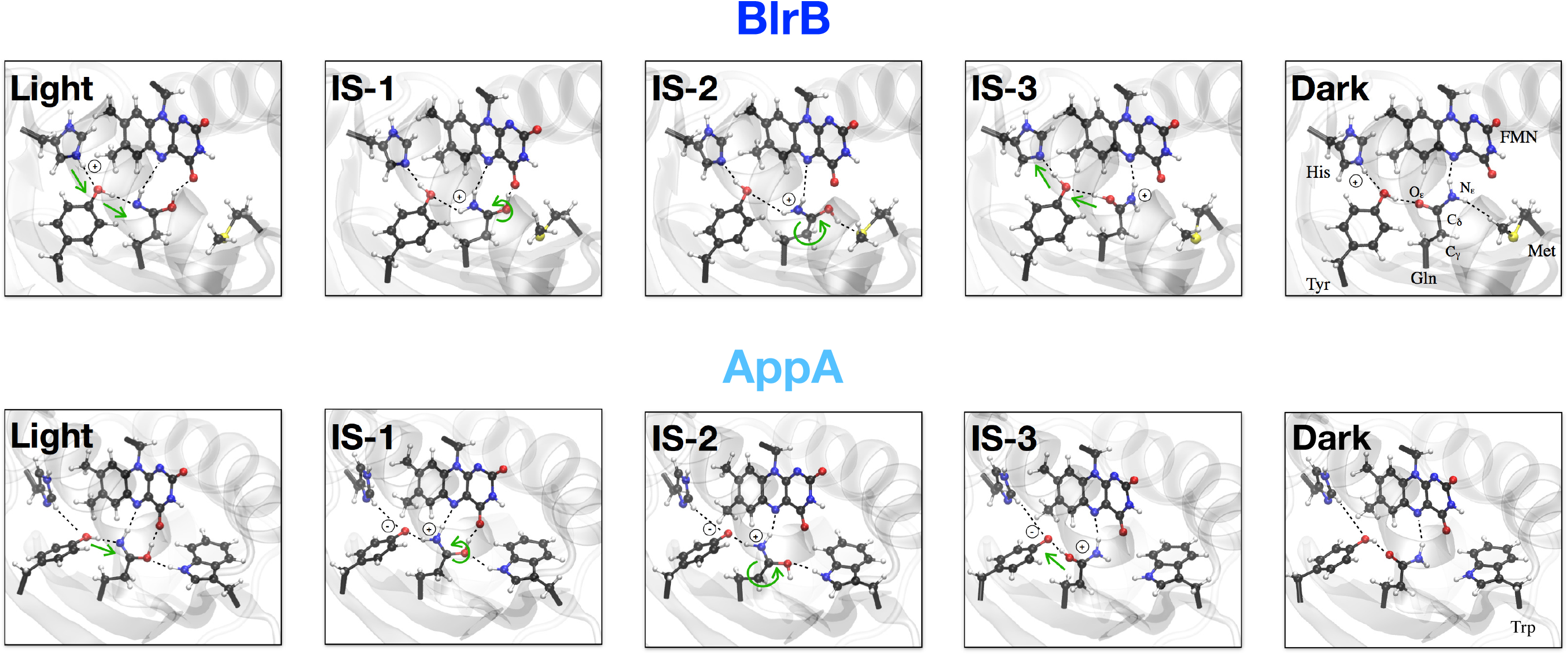
Light- to dark state transition mechanism, with three labelled intermediate states (IS) in the BlrB and AppA BLUF proteins, obtained from the path-steered MD simulations. The mechanism in BlrB is in agreement with that proposed by Khrenova and coworkers [42]. The AppA mechanism shows several modifications, including a neutral His and Tyr being the first proton donor.

Here, we investigate the dark-state recovery of the BlrB and the AppA proteins. Given the current consensus on the Gln rotation and tautomerization [11, 36], we initially assume a mechanism similar to the one proposed by Khrenova and collaborators for the BlrB protein [42]. For the AppA protein, we consider the Tyr—rather than a protonated His—as the first proton donor. This consideration follows experimental measurements showing that AppA has a much greater sensitivity to Tyr triple-fluorination; decreasing its recovery time 4000-fold [16]. Additionally, for AppA, we take into account the switch from the Trp_in_/Met_out_ (light) to the Trp_out_/Met_in_ (dark) conformation [41]. We present simulations at the QM/MM level for the hydrogen-bond rearrangement near the chromophore, for both BlrB and AppA, and at the MM level for the Trp/Met switching in AppA. Moreover, rather than obtaining potential-energy profiles along energy-minimized intermediate states, we perform QM/MM-based molecular dynamics (MD) to validate our mechanistic pathways and calculate free-energy profiles. This task, which is particularly expensive for QM/MM MD, becomes feasible thanks to our in-house developed path-based free-energy methods [9, 59, 60]. By sampling along an adaptive light-to-dark path in the space of key transition descriptors, i.e. collective variables (CVs), we efficiently calculate a free-energy profile. For our QM/MM simulations of the change in hydrogen-bonding within the active site, we use CVs inspired by the currently accepted mechanism, and we render free-energy profiles for the AppA and BlrB proteins, which we compare with previous work and experimental recovery rates [76, 16, 17].

## 2 Simulation protocol

### 2.1 Molecular mechanics

System preparation is done using GROMACS 2018.8 [2, 72]. For BlrB, we start with the X-ray structure 2BYC (chain A) [39] from the Protein Data Bank [3]. Same as in [42], we consider Lys and Arg residues to be positively charged, and Glu and Asp residues to be negatively charged. Since His73 is considered to be the first proton donor, we rotate it and protonate it at both nitrogen atoms in order to facilitate a proton transfer to Tyr9, as done in [42]. All other residues are protonated according to default GROMACS protocols. The Gln is in amide form—for which force-field parameters are readily available—with its oxygen pointing toward the Tyr, which corresponds to the dark state. The system is centered in a cubic box with a distance of 1 nm between the protein and the edge of the box. The protein is modeled with the CHARMM36 force field [33]. A flavin mononucleotide (FMN), which substitutes the FAD, is modelled with parameters from [13]. We solvate the system in TIP3P water [38] and then neutralize the charge by adding 150 mM NaCl [23]. The system is then energy-minimized. We use the canonical sampling via velocity rescaling (CSVR) thermostat [5]—set at 300 K with a time constant of 0.1 ps—and the Parrinello-Rahman barostat [57]—set at 1 bar with a time constant of 1 ps. The LINCS algorithm [31] was used to constrain bonds involving hydrogen atoms. With a step of 2 fs, we perform 100 ns of equilibration.

For AppA, we start with the PDB X-ray structure 1YRX (chain A) [1], which has the Trp_in_/Met_out_ conformation assigned to the light state. The Gln is also in the light-state orientation, i.e. with its oxygen pointing toward the C_4_ O carbonyl of the FAD, but it is in amide form, due to force-field parameter availability. We consider all His residues protonated only at their N_*ε*_ atoms; including the His85 near the FMN, since Tyr21 is now considered the first proton donor. The rest of the preparation is done in the same manner as for BlrB.

### 2.2 Quantum mechanics/molecular mechanics

We start from the final structures from the force-field MD equilibration runs described above. We run QM/MM MD using CP2K [34, 47]. For BlrB, following the example of Khrenova and coworkers [42], we include in the QM zone the residues Tyr9, Ser11, Asn33, Gln51, His73, Met94 and the FMN, which have a total of 92 atoms. The QM box size is of 25 *×* 25 *×* 25 Å. We use the Perdew-Burke-Ernzerhof (PBE) exchange-correlation functional [58]—which has a very favorable accuracy versus computational cost performance—together with Grimme’s D3 dispersion correction [24]. We use Goedecker-Teter-Hutter (GTH) pseudopotentials [18, 26] to represent valence-core interactions, and a double-zeta valence with polarization (DZVP-GTH-PBE) basis set with a plane wave cutoff of 400 Ry. All QM residues are capped by hydrogen atoms, saturating the C_*α*_ with a radius of 1.4 Å and a force scaling factor of 1.5. The QM region is embedded electrostatically inside the MM environment, which is handled with the same force fields as in the MM protocol. After a geometry optimization, we equilibrate for 5 ps with a time step of 0.5 fs. We use the CSVR thermostat [5] set at 300 K with a time constant of 0.5 ps, and with a thermal region for the QM atoms. The time evolution of the root-mean-square deviation (RMSD) of the QM atom positions plateaus at ∼ 1 Å after 2 ps; and the energy drift is *<*0.4 K/ps per atom; indicating a stable setup. A smaller QM zone was tested—only with Tyr9, Gln51, His73 and the FMN—but dismissed after noticing lesser stability, i.e. the RMSD time evolution not stabilizing after 5 ps and an energy drift of ∼ 0.6 K/ps per atom. After the dark-state is equilibrated, we use path-steered MD—with parameters similar to the ones described in the next section—to rotate and tautomerize the Gln. Then, we re-equilibrate at the light-state.

For the AppA protein, we repeat a similar protocol, including in the QM zone the residues Tyr21, Ser23, Asn45, Gln63, His85, Trp104 and the FMN, which account for a total of 98 atoms. We use the same QM/MM parameters as for BlrB. We manually change the Gln to the ZZ-imide form, before equilibrating in the light state. To obtain the dark state structure, we again use path-steered MD with settings similar to the ones described below, and then re-equilibrate.

### 2.3 Path-constrained molecular dynamics

Molecular transitions that require crossing over a free-energy barrier, such as chemical reactions or conformational changes, are typically rare events in the timescales accessible to MD simulations; specially at higher levels of theory like QM/MM. Standard free-energy methods, such as constrained MD, steered MD or metadynamics, can sample these rare events by exerting a biasing potential on a small set of CVs and return an insightful free-energy surface. However, the convergence time of these calculations suffers from exponential scaling with respect to the number of CVs; meaning that complex transitions that involve many CVs are often unfeasible. Here, we employ the path-based that was first introduced in [9] and extended in [59]. In short, rather than sampling the entire CV-space, we sample the one-dimensional progress component, *s*, along an adaptive path connecting two known stable states. Most standard free-energy methods—such as constrained MD [6, 8] or steered MD [25, 37]—can be directly used along the path, and provide valuable mechanistic details, as exemplified in [60].

For the BlrB BLUF protein, we use eight CVs to describe the tautomerization and rotation of the Gln (see Fig. 3):

- *c*_Tyr-H1_: coordination number^1^ of the Tyr O with the proton H1^2^.
- *c*_His-H1_: coordination number of the His N_*ε*_ with the proton H1.
- *c*_Gln-H2_: coordination number of the Gln N_*ε*_ with the proton H2.
- *c*_Tyr-H2_: coordination number of the Tyr O with the proton H2.
- *t*_O-H_: dihedral angle formed by the C_*γ*_ -C_*δ*_ -O_*ε*_ -H3. This angle is shifted by +5.5 rad in order to capture the desired direction of rotation, i.e. with the O-H pointing toward the FMN at the mid-rotation.
- *t*_Gln_: pseudo-dihedral angle formed by: the axis between the C_*γ*_ and the C_*δ*_ of Gln; the first vector from the C_*α*_ of the previous residue of Gln to C_*α*_ of the following residue; and the second vector from the N_*ε*_ to the O_*ε*_ of Gln. The direction of rotation is such that the Gln O points toward the FMN at the mid-rotation.
- *c*_Tyr-H3_: coordination number of the Tyr O with the proton H3.
- *c*_Gln-H3_: coordination number of the Gln O_*ε*_ with the proton H3.

The initial and final points of the light-to-dark path are based on the average values of the CVs at each of the stable states (see Table 1). We normalize the CVs such that their values are 1 at the light state, and 0 at the dark state^3^. The path has 20 nodes, with 10 extra trailing nodes at the beginning and at the end. The light state corresponds to path progress component *s* = 1.0, and the dark state to *s* = 0.0. We shape the path in four steps, of ∼ 5 nodes each, according to the steps proposed by Khrenova and coworkers [42]. The eight CVs decrease from 1 to 0 on the following order: First, *c*_Tyr-H1_, *c*_His-H1_, *c*_Gln-H2_ and *c*_Tyr-H2_; second, *t*_O-H_; third, *t*_Gln_; and fourth, *c*_Tyr-H3_ and *c*_Gln-H3_, while *c*_Tyr-H1_ and *c*_His-H1_ increase again from 0 to 1. Given the high computational cost of QM/MM MD, and the fact a proposed mechanism is already available, we do not adapt the path on the fly. Instead, we allow the sampling to moderately deviate from the assumed pathway. We do this by setting a “tube” potential—i.e. a harmonic restraint on the distance component from the path, *z*—with an offset of 0.5 normalized units, and a force constant of 1,000 kcal/mol. This implies that the sampling of the transition may drift up to 0.5 units from the path in the normalized CV-space.

**Table 1:** Average values of the eight CVs during the QM/MM light- and dark-state 5 ps equilibrations of the BlrB BLUF protein.

| state / CV | $c_{\text{Tyr-H1}}$ | $c_{\text{His-H1}}$ | $c_{\text{Gln-H2}}$ | $c_{\text{Tyr-H2}}$ | $t_{\text{O-H}}$ | $t_{\text{Gln}}$ | $c_{\text{Tyr-H3}}$ | $c_{\text{Gln-H3}}$ |
| --- | --- | --- | --- | --- | --- | --- | --- | --- |
| Light | 0.3 | 0.9 | 0.3 | 0.9 | -2.2 | 2.5 | 0.0 | 0.9 |
| Dark | 0.3 | 0.9 | 0.9 | 0.0 | 1.2 | -1.8 | 0.9 | 0.3 |

To obtain free-energy profiles, we perform path-constrained MD, with samples from *s* = 0.0 to *s* = 1.0, every 0.05 path progress units. We use stiff harmonic restraints, with a force constant of 100,000 kcal/mol. The large value of the force constant is due to the normalization of the path progress, on top of the normalization of the CVs. The samples are moved to the desired positions via path-steered MD, starting from *s* = 1.0, with a speed of 2.0 path progress units per ps. The sample at *s* = 1.0 is steered from *s* = 0.0, to verify the reversibility of the transition. After being steered to their respective positions, we let the systems equilibrate for at least 0.5 ps before recording forces for 1 ps. For the samples between *s* = 0.35 to *s* = 0.55, which involve the slow rotation of the Gln, we double the equilibration and sampling time. Finally, we obtain free-energy profiles via numerical integration of the average forces, and cubic spline interpolation.

Some restraints are included in the simulations, all with force constants of 100 kcal mol^*−*1^ Å ^*−*2^. The distance between the Tyr O and the second proton of the Gln N_*ε*_ in the dark-state is restrained to be larger than 2.5 Å to prevent proton transfer. The distance between the Tyr O and the Gln C_*δ*_ is restrained to be larger than 2.5 Å to avoid a reaction with the mid-rotated Gln. The distance between the His N_*ε*_ and the Tyr O is restrained at 2.8 Å to ensure its favorable position for the first concerted proton transfer. The tendency of the His flipping out of the active site and into the solvent, greater in MM than in QM/MM MD, has previously been observed in [42].

For the AppA BLUF protein we repeat the same protocol, with a few adjustments. Because we do not consider a proton transfer from the His (see Fig. 3), the CVs *c*_Tyr-H1_ and *c*_His-H1_, as well as the restraint between the His N_*ε*_ and the Tyr O, are not used. To prevent proton transfer back to the Tyr during the O-H and Gln rotation steps, the offset of the tube potential is decreased to 0.3 normalized units. For the samples at *s* = 0.65, *s* = 0.60 and *s* = 0.45 the offset had to be further decreased to 0.2 normalized units; and for the sample at *s* = 0.35, to 0.1 normalized units. The sample at *s* = 0.7 is the only one with an offset of 0.5 normalized units.

## 3 Results and discussion

### 3.1 Light- and dark-state structures

In Table 2 we show the hydrogen-bond occupancies for the BlrB and AppA BLUF proteins, as measured during the QM/MM MD simulations in the light and dark states. The hydrogen bonds are illustrated with dashed lines in Fig. 3. In general, we observe good agreement with the currently accepted structures [11, 36] and with previous QM/MM simulations [20].

**Table 2:** Hydrogen-bond occupancies during the QM/MM MD equilibrations of the BlrB and AppA BLUF proteins in the light and dark states. A hydrogen bond is considered formed if the donor-acceptor distance is *<*3.2 Å. We evaluate the distances every 50 fs for the 5 ps trajectories.

| H-bond | BlrB Light | BlrB Dark | AppA Light | AppA Dark |
| --- | --- | --- | --- | --- |
| Gln $O_\epsilon$-FMN $O_4$ | 78% | 0% | 100% | 3% |
| Gln $N_\epsilon$-FMN $N_5$ | 4% | 48% | 72% | 12% |
| Gln $O_\epsilon$-Met S/Trp N | 3% | 0% | 84% | 0% |
| Gln $N_\epsilon$-Tyr O | 93% | 0% | 100% | 0% |
| Gln $N_\epsilon$-FMN $O_4$ | 0% | 65% | 0% | 91% |
| Gln $N_\epsilon$-Met S/Trp N | 0% | 0% | 0% | 7% |
| Gln $O_\epsilon$-Tyr O | 0% | 100% | 0% | 94% |

In the light state of both proteins, the Gln O_*ε*_ makes a persistent hydrogen bond with the O_4_ of the FMN; agreeing with previous spectroscopic analyses [48]. The Gln O_*ε*_ -FMN O_4_ hydrogen bond appears to be stronger in AppA (100% occupancy) than in BlrB (78%). The second Gln-to-FMN hydrogen bond, which involves the N_*ε*_ and N_5_ atoms, is much more persistent in AppA (72%) than in BlrB (4%). Moreover, the interaction of the Gln O_*ε*_ with Trp in AppA (84%) is much stronger than with the Met in BlrB (3%) These stable-state trends already show consistency with the longer recovery time in the AppA protein, where it is likely more difficult to break the hydrogen bonds of the Gln with the FMN and the Trp. Both proteins have persistent hydrogen-bonding between the Gln N_*ε*_ and Tyr O.

In the dark state, we also see some differences between the two BLUF proteins. The Gln N_*ε*_ of BlrB makes hydrogen bonds with both the FMN N_5_ (48%) and the FMN O_4_ (65%). On the other hand the Gln N_*ε*_ of AppA is significantly more hydrogen-bonded to the FMN O_4_ (91%) than to the FMN N_5_ (12%). This is due to differences in the alignment of the ZZ-imide Gln. In both proteins, the interaction of the Gln N_*ε*_ with either the Trp or the Met is very weak. The hydrogen-bonding of the Gln O_*ε*_ with the Tyr O is persistent in both proteins.

### 3.2 Glutamine tautomerization and rotation

In Fig. 4 we show free-energy profiles for the light-to-dark hydrogen-bond change resolved with QM/MM-based path-constrained MD. For the BlrB protein, the first barrier—from *s* ≈ 1.0 to *s* ≈ 0.7—corresponds to the first concerted proton transfers from the His to the Tyr, and from the Tyr to the Gln. The second barrier, *s* ≈ 0.7 to *s* ≈ 0.5, tracks the O-H rotation, while the third barrier, *s* ≈ 0.5 to *s* ≈ 0.3, does so for the Gln rotation. Finally, the last barrier—from *s* ≈ 0.3 to *s* ≈ 0.0—represents the concerted proton transfers from the Gln to the Tyr, and from the Tyr to the His (see Fig. 3).

**Figure 4:**
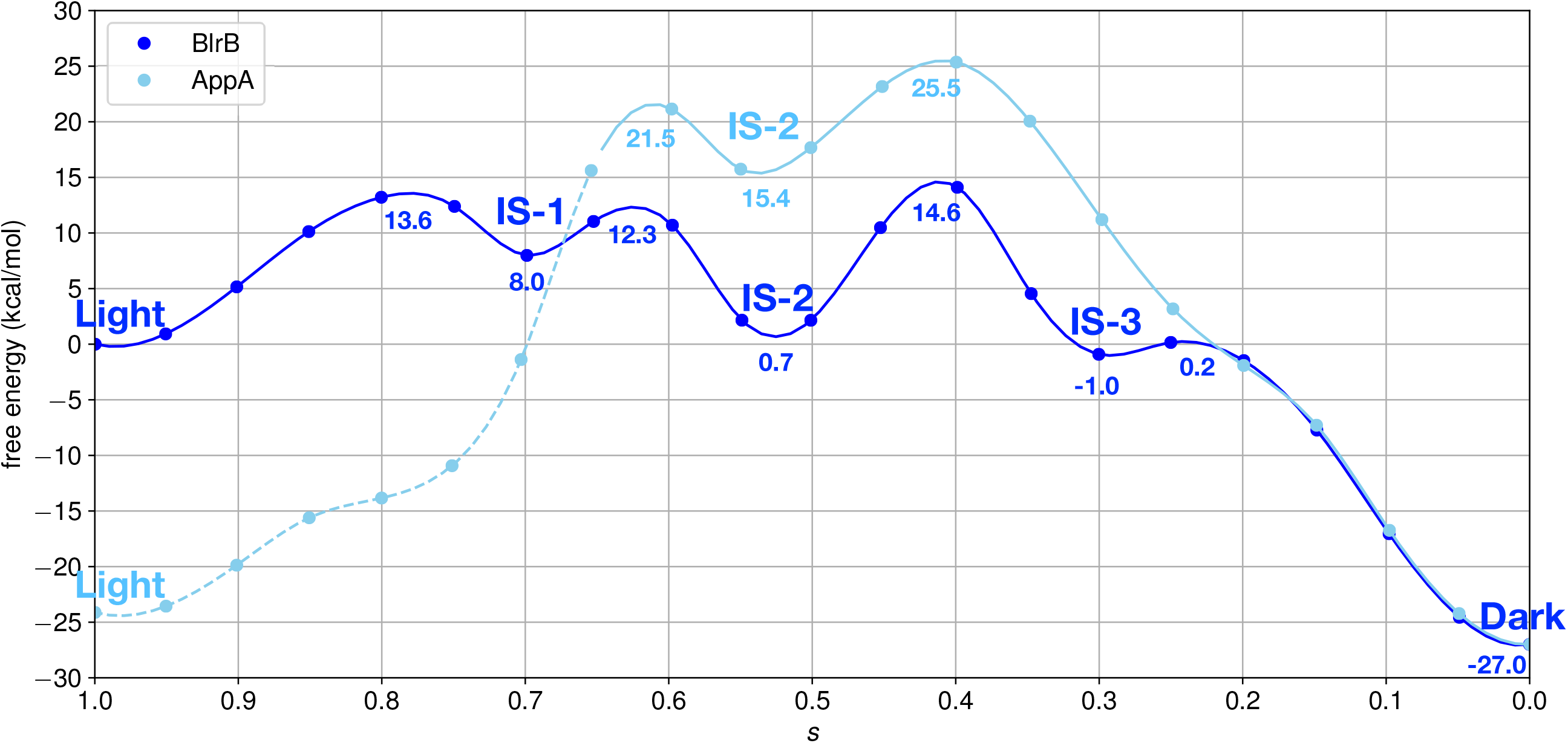
Light- to dark-state free-energy profiles of the BlrB and AppA BLUF proteins obtained by path-constrained MD with steps and intermediate states (IS) as described in Fig. 3. The free-energy obtained from numerical integration of the constraining forces is shown with solid points, and a cubic spline interpolation is shown with lines. A dashed line shows the section of the AppA BLUF free-energy profile that corresponds to the first Tyr-to-Gln proton transfer.

Our result for the BlrB protein is consistent with the previously published potential-energy profile [42], especially regarding the O-H rotation and Gln rotation barriers. The Gln rotation is the rate-determining step, for which we calculate a barrier of 14.6 kcal/mol, in perfect agreement with the previous result. The experimental recovery time of 2 s [76] implies a rate-determining barrier of ∼ 18 kcal/mol ^4^. The mismatch of ∼ 3.4 kcal/mol between the calculations and the experiments could be attributed to the size of the QM region, or other QM and MM parameters. Continuing our comparison to the potential-energy calculation, we calculate a much higher barrier (13.6 kcal/mol) for the first proton transfers than previously reported (2.2 kcal/mol). The increased barrier might be due to the assumption that the His-to-Tyr and Tyr-to-Gln proton transfers are concerted, which comes from the single transition state reported in [42]. In Fig. 5—which shows the sampling projected on each CV—we observe that, during the first proton transfers, *c*_His-H1_ and *c*_Tyr-H2_ deviate from the path by an amount close to the limit of 0.5 normalized units; indicating that the proton transfers might indeed occur in a stepwise manner. We also record a positive free-energy difference (8.0 kcal/mol) for the first proton transfers, opposite to the previous potential-energy study (−2.1 kcal/mol); pointing to less likely excursions to the first intermediate. The disagreement might also be due to differences in the models, such as the choice for the DFT exchange-correlation functional, which is PBE in this work versus PBE0 in the former work.

**Figure 5:**
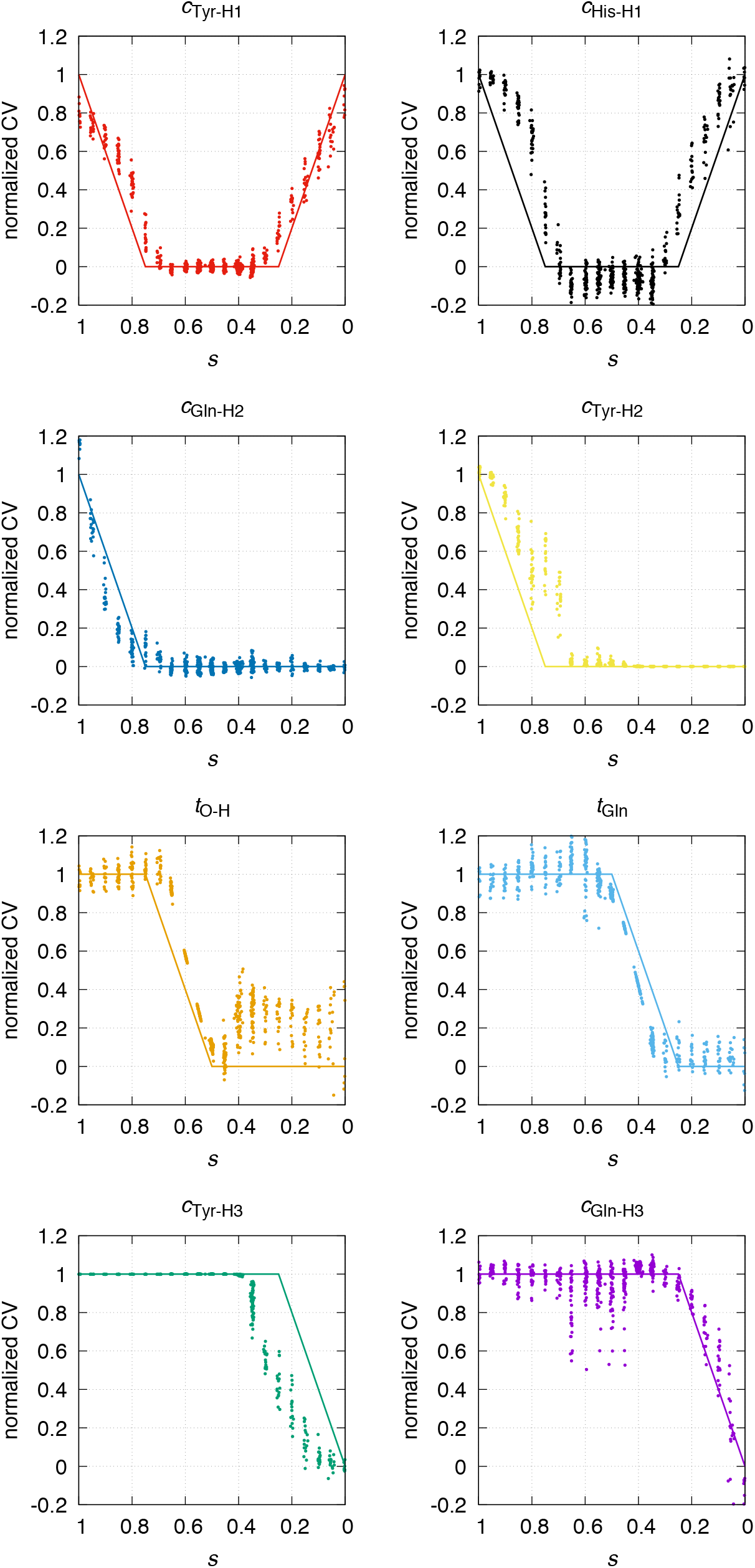
Path-constrained MD sampling projected onto each of the eight CVs and the path progress, *s*, during the calculation for the dark-state recovery of the BlrB BLUF protein.

To analyze the mechanism, in Fig. 5 we show the path and the sampling projected onto each of the eight CVs. For most cases, the sampling remains close to the path. As mentioned before, the largest deviations from the path occur for the concerted proton transfers. While it is common that the coordination numbers of the acceptors change before those of the donors, the deviations indicate that the His-to-Tyr and Tyr-to-Gln proton transfers might be more favored when occurring in a stepwise manner. The coordination number *c*_Gln-H3_ shows deviations in the range from *s* = 0.7 to *s* = 0.5, which are caused by the FMN C_4_ O attracting the proton during the rotation of the O-H group. Also, the torsion *t*_O-H_ shows deviations near the dark state, which we observe in the corresponding equilibration as well, because the proton is transferred to the Tyr and no longer forms part of the O-H group.

In Fig. 4, we also show a free-energy profile for the recovery of the AppA BLUF protein. The free energy of the Tyr-to-Gln proton transfer, taking place from *s* ≈ 1.0 to *s* ≈ 0.7, is raised by almost ∼ 25 kcal/mol due to the lack of a proton donor for the Tyr. Moreover, the first and third intermediate states (IS-1 and IS-3), as shown in Fig. 3, do not show any metastability. The free-energy profile strongly suggests that the AppA protein, like BlrB, requires a residue to donate a proton to the Tyr. To explain the difference in recovery times—from 2 s in BlrB to ∼25 min in AppA—we propose the role of the Trp_in_/Met_out_ conformation. To compare the free-energy profiles of both proteins in Fig. 4, we assume that the Tyr deprotonation does not affect the AppA free-energy calculation beyond the range from *s* ≈ 1.0 to *s* ≈ 0.7. In BlrB, starting from the top of the second barrier (12.3 kcal/mol), which corresponds to the O-H rotation, the second intermediate state (IS-2) implies a lowering of 11.6 kcal/mol in free energy. In AppA, IS-2 is only 6.1 kcal/mol lower than the peak of second barrier. This increased metastability of IS-2 in BlrB is likely caused by the hydrogen bond of the Gln O-H group with the Met (98% occupancy), which is replaced by the Trp in AppA. Moreover, the third barrier is only 2.3 kcal/mol higher than the second barrier in BlrB, while in AppA it is 4.0 kcal/mol higher; indicating that the Trp hinders Gln rotation; agreeing with the stronger hydrogen-bonding of the light-state Gln O_*ε*_ (84%) shown in Table 1. Under the reasonable assumption that introducing a protonated His in AppA would level the two-free energy profiles at the top of the second barrier (see dotted line in Fig. 4), then the barrier of the rate limiting step—i.e. the Gln rotation—of AppA would be of ∼ 16.3 kcal/mol. While this number is still lower than the ∼ 22 kcal/mol that might be expected from the ∼ 25 min recovery time of AppA, this would mean that both free-energy calculations underestimate the barrier by a similar factor: 0.8 in BlrB and 0.7 in AppA.

The mechanistic details of the AppA recovery, shown in Fig. 6, are similar to those of BlrB. Again, the coordination numbers deviate slightly for the path, with the proton acceptors changing before the donors. Same as in BlrB, the coordination number *c*_Gln-H3_ fluctuates from *s* = 0.7 to *s* = 0.5, because the C_4_ O hinders the rotation of the O-H group, and the torsion *t*_O-H_ shows fluctuations near the dark state, also seen in the equilibration. The coordination number *c*_Tyr-H2_ shows larger fluctuations than in BlrB from *s* = 0.7 to *s* = 0.4, because the deprotonated Tyr is still interacting with the proton at the Gln C_*ε*_.

**Figure 6:**
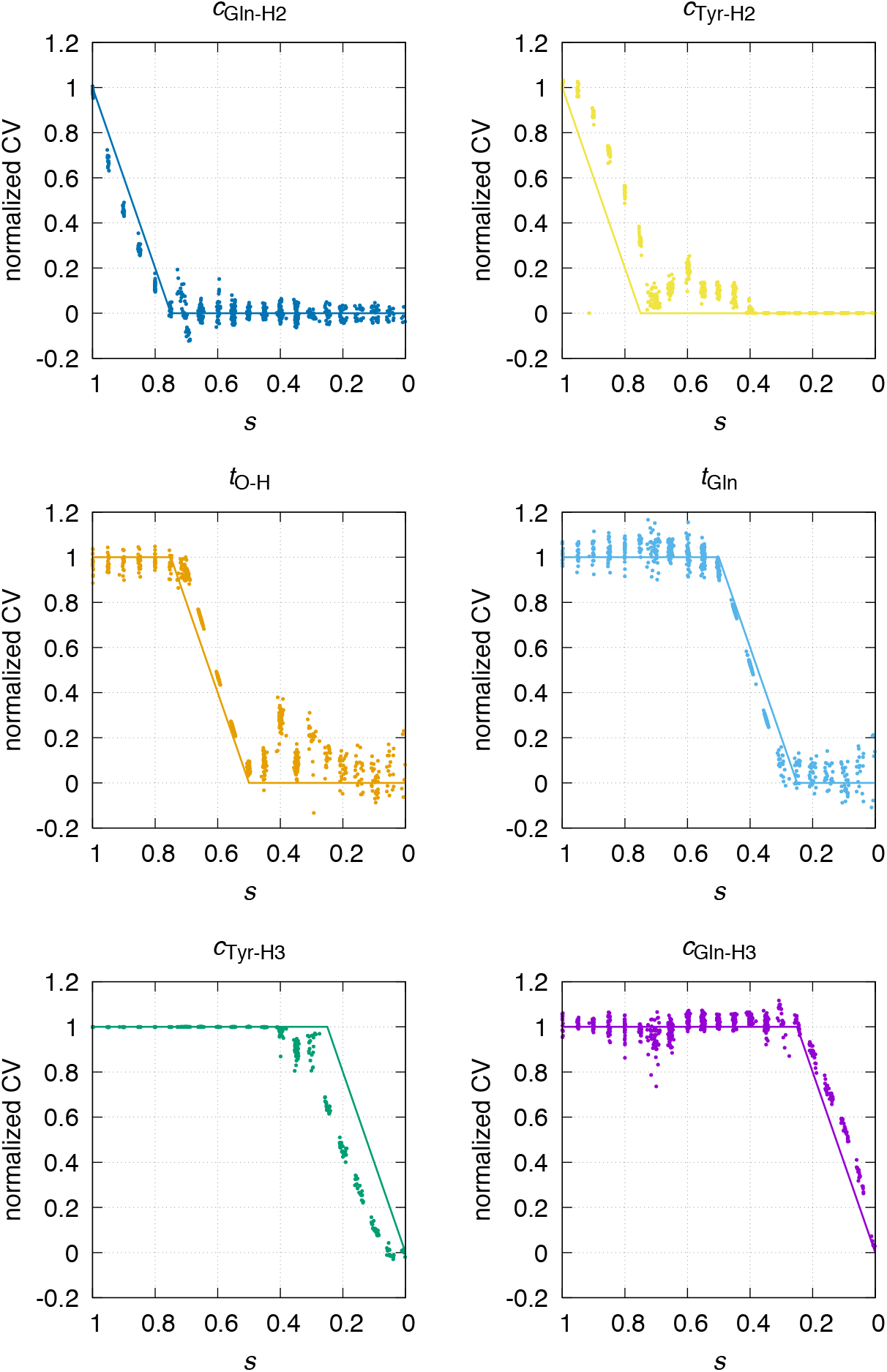
Path-constrained MD sampling projected onto each of the six CVs and the path progress, *s*, during the calculation for the dark-state recovery of the AppA BLUF protein.

## 4 Conclusion

For the first time, dynamics and free-energy calculations where performed of the Gln tautomerization and rotation associated with the deactivation of BLUF photoreceptors. In a QM/MM MD setup, we corroborate the stability of the currently accepted light- and dark-state structures in two different BLUF proteins: BlrB and AppA. We also confirm the BlrB dark-state recovery mechanism proposed by Khrenova and coworkers [42] via path-steered and -constrained MD; obtaining a remarkable agreement in the energy barriers. Said mechanism involves several steps—i.e. proton transfers between a protonated His, a conserved Tyr and the Gln, as well as rotations of the Gln—whose handling is streamlined in path-based calculations.

Aided by the efficiency of path-based calculations, we also obtain a free-energy profile for the dark-state recovery of the AppA protein. Based on the reported high sensitivity to Tyr-fluorination of this protein, we consider its Tyr as a first proton donor, rather than the protonated His considered in BlrB. However, our calculated free-energy profile strongly points that the AppA protein, same as BlrB, must have an initial proton donor different from the Tyr. We propose, based on our free-energy profiles, that the ∼ 750-fold longer recovery time of AppA with respect to BlrB is due to the Trp_in_/Met_out_ conformation of the former. The respective interactions of the Met or Trp modulate the barrier for the Gln rotation, i.e. the rate-determining step of the recovery. In AppA, it has already been observed that the Trp flips from inside to outside of the active site as the protein recovers the dark state. The exact concertedness of the Trp/Met switch with the Gln tautomerization and rotation might determine the exact recovery rate. Larger, or adaptive QM regions may help including the Trp/Met conformational change in the same simulation as the Gln rotation and tautomerization.

Our scheme for QM/MM path-based sampling enables future calculations in BLUF proteins or similar systems undergoing intricate hydrogen-bond rearrangements. This includes a calculation of the AppA dark-state recovery with a protonated His, as well as fluorinated Tyr mutants, which can be compared to experiments. Robust multiscale schemes and sampling methods hold the key to further understand the variations of signaling times across BLUF proteins, and to predict directed mutations to control their behavior in optogenetic or biosensor applications.

## Acknowledgements

The work has been performed under the Project HPC-EUROPA3 (INFRAIA-2016-1-730897), with the support of the EC Research Innovation Action under the H2020 Programme; in particular, the author gratefully acknowledges the support of Prof. Carme Rovira (Departament de Química Inorgànica i Orgànica & Institut de Química Teórica i Computacional, Universitat de Barcelona) and the computer resources and technical support provided by the Barcelona Supercomputing Center.A.P.A.O. received funding from the Mexican National Council for Science and Technology (CONACYT). A.P.A.O. acknowledges I. Schapiro, M.-A. Mroginski and Y. Yogev for organizing the inspiring CECAM workshop “Frontiers in Multiscale Modelling of Photoreceptor Proteins”.

## Data availability

All the data and PLUMED input files required to reproduce the results reported in this paper will be made available on PLUMED-NEST (www.plumed-nest.org), the public repository of the PLUMED consortium [4].

## A Appendix

### A.1 Path-constrained molecular dynamics sampling

Fig. 5 and Fig. 6 show the path-constrained MD sampling of the dark-state recovery mechanism in the BlrB and the AppA BLUF proteins, respectively. The sampling remains mostly close to the path for both cases. The deviations are discussed in the main text.

## Footnotes

1 Coordination numbers are calculated as *c*_*i j*_ = [1 − (*r*_*i j*_*/r*_0_)^6^]*/*[1 − (*r*_*i j*_*/r*_0_)^12^], where *r*_*i j*_ is the distance between the two atoms involved and *r*_0_ = 1.5 Å.

2 We label protons based on their positions in the light state (see Fig. 3): H1 is the proton of the His N_*ε*_, H2 is the proton of the Tyr O, and H3 is the proton of the Gln O_*ε*_.

3 The CV normalization is done with *x*_norm_ = (*x−x*_dark_)*/*(*x*_light_ *−x*_dark_), where *x* is the corresponding CV.

4 Assuming a free energy barrier Δ*F*^*\**^ = *N*_A_*k*_B_*T* ln[(*kh*)*/*(*k*_B_*T*)], with the Avogadro constant *N*_A_, Boltzmann constant *k*_B_, Planck constant *h*, rate constant *k* and temperature *T*.

## References

[1] Spencer Anderson, Vladimira Dragnea, Shinji Masuda, Joel Ybe, Keith Moffat, and Carl Bauer. Structure of a novel photoreceptor, the BLUF domain of AppA from Rhodobacter sphaeroides. Biochemistry, 44(22):7998–8005, 2005.

[2] Herman J C Berendsen, David van der Spoel, and Rudi van Drunen. GROMACS: a message-passing parallel molecular dynamics implementation. Comput. Phys. Commun., 91(1-3):43–56, 1995.

[3] H. M. Berman, J. Westbrook, Z. Feng, G. Gilliland, T. N. Bhat, H. Weissig, I. N. Shindyalov, and P. E. Bourne. The Protein Data Bank. Nucleic Acids Res., 28:235–242, 2000.

[4] Massimiliano Bonomi, Giovanni Bussi, Carlo Camilloni, Gareth A Tribello, Pavel Banáš, Alessandro Barducci, Mattia Bernetti, Peter G Bolhuis, Sandro Bottaro, Davide Branduardi, et al. Promoting transparency and reproducibility in enhanced molecular simulations. Nat. Methods, 16(8):670–673, 2019.

[5] G. Bussi, D. Donadio, and M. Parrinello. Canonical sampling through velocity rescaling. J. Chem. Phys., 126:014101, 2007.

[6] E. A. Carter, G. Ciccotti, J. T. Hynes, and R. Kapral. Constrained reaction coordinate dynamics for the simulation of rare events. Chem. Phys. Lett, 156:472, 1989.

[7] John M Christie, Jayde Gawthorne, Gillian Young, Niall J Fraser, and Andrew J Roe. LOV to BLUF: flavoprotein contributions to the optogenetic toolkit. Mol. Plant., 5(3):533–544, 2012.

[8] W. K. den Otter and W. J. Briels. The calculation of free-energy differences by constrained molecular dynamics simulations. J. Chem. Phys., 109:4139, 1998.

[9] Grisell Díaz Leines and Bernd Ensing. Path finding on high-dimensional free energy landscapes. Phys. Rev. Lett, 109(2):020601, 2012.

[10] Tatiana Domratcheva, Bella L Grigorenko, Ilme Schlichting, and Alexander V Nemukhin. Molecular models predict light-induced glutamine tautomerization in BLUF photoreceptors. Biophys. J., 94(10):3872–3879, 2008.

[11] Tatiana Domratcheva, Elisabeth Hartmann, Ilme Schlichting, and Tilman Kottke. Evidence for tautomerisation of glutamine in BLUF blue light receptors by vibrational spectroscopy and computational chemistry. Sci. Rep., 6:22669, 2016.

[12] Martin J Field, Paul A Bash, and Martin Karplus. A combined quantum mechanical and molecular mechanical potential for molecular dynamics simulations. J. Comput. Chem, 11(6):700–733, 1990.

[13] Peter L Freddolino, Kevin H Gardner, and Klaus Schulten. Signaling mechanisms of lov domains: new insights from molecular dynamics studies. Photochem. Photobiol. Sci., 12(7):1158–1170, 2013.

[14] Tomotsumi Fujisawa and Shinji Masuda. Light-induced chromophore and protein responses and mechanical signal transduction of BLUF proteins. Biophys. Rev., 10(2):327–337, 2018.

[15] Tomotsumi Fujisawa, Shinji Masuda, Satoshi Takeuchi, and Tahei Tahara. Femtosecond time-resolved absorption study of signaling state of a bluf protein pixd from the cyanobacterium synechocystis: Hydrogen-bond rearrangement completes during forward proton-coupled electron transfer. J. Phys. Chem. B, 125(44):12154–12165, 2021.

[16] Agnieszka A Gil, Allison Haigney, Sergey P Laptenok, Richard Brust, Andras Lukacs, James N Iuliano, Jessica Jeng, Eduard H Melief, Rui-Kun Zhao, EunBin Yoon, et al. Mechanism of the AppABLUF Photocycle Probed by Site-Specific Incorporation of Fluorotyrosine Residues: Effect of the Y21 p K a on the Forward and Reverse Ground-State Reactions. J. Am. Chem. Soc., 138(3):926–935, 2016.

[17] Agnieszka A Gil, Sergey P Laptenok, James N Iuliano, Andras Lukacs, Anil Verma, Christopher R Hall, Grace E Yoon, Richard Brust, Gregory M Greetham, Michael Towrie, et al. Photoactivation of the BLUF protein PixD probed by the site-specific incorporation of fluorotyrosine residues. J. Am. Chem. Soc., 139(41):14638–14648, 2017.

[18] S Goedecker, M Teter, and Jürg Hutter. Separable dual-space gaussian pseudopotentials. Phys. Rev. B, 54(3):1703, 1996.

[19] Joshua J Goings and Sharon Hammes-Schiffer. Early Photocycle of Slr1694 Blue-Light Using Flavin Photoreceptor Unraveled through Adiabatic Excited-State Quantum Mechanical/Molecular Mechanical Dynamics. J. Am. Chem. Soc., 141(51):20470–20479, 2019.

[20] Joshua J Goings, Pengfei Li, Qiwen Zhu, and Sharon Hammes-Schiffer. Formation of an unusual glutamine tautomer in a blue light using flavin photocycle characterizes the light-adapted state. Proc. Natl. Acad. Sci. USA, 117(43):26626–26632, 2020.

[21] Joshua J Goings, Clorice R Reinhardt, and Sharon Hammes-Schiffer. Propensity for proton relay and electrostatic impact of protein reorganization in Slr1694 BLUF photoreceptor. J. Am. Chem. Soc., 140(45):15241–15251, 2018.

[22] Mark Gomelsky and Gabriele Klug. BLUF: a novel FAD-binding domain involved in sensory transduction in microorganisms. Trends Biochem. Sci., 27(10):497–500, 2002.

[23] Puja Goyal and Sharon Hammes-Schiffer. Role of active site conformational changes in photocycle activation of the AppA BLUF photoreceptor. Proc. Natl. Acad. Sci. USA, 114(7):1480–1485, 2017.

[24] Stefan Grimme, Jens Antony, Stephan Ehrlich, and Helge Krieg. A consistent and accurate ab initio parametrization of density functional dispersion correction (DFT-D) for the 94 elements H-Pu. J. Chem. Phys., 132(15):154104, 2010.

[25] Helmut Grubmüller, Berthold Heymann, and Paul Tavan. Ligand binding: molecular mechanics calculation of the streptavidin-biotin rupture force. Science, 271(5251):997–999, 1996.

[26] C Hartwigsen, Sephen Gœdecker, and Jürg Hutter. Relativistic separable dual-space gaussian pseudopotentials from h to rn. Phys. Rev. B, 58(7):3641, 1998.

[27] Shaima Hashem, Giovanni Battista Alteri, Lorenzo Cupellini, and Benedetta Mennucci. Integrated computational study of the light-activated structure of the appa bluf domain and its spectral signatures. J. Phys. Chem. A, 2023.

[28] Shaima Hashem, Lorenzo Cupellini, Filippo Lipparini, and Benedetta Mennucci. A polarisable qm/mm description of nmr chemical shifts of a photoreceptor protein. Mol. Phys., 118(19-20):e1771449, 2020.

[29] Shaima Hashem, Veronica Macaluso, Michele Nottoli, Filippo Lipparini, Lorenzo Cupellini, and Benedetta Mennucci. From crystallographic data to the solution structure of photoreceptors: the case of the appa bluf domain. Chem. Sci., 12(40):13331–13342, 2021.

[30] YongLe He, Agnieszka A Gil, Sergey P Laptenok, Anam Fatima, Jinnette Tolentino Collado, James N Iuliano, Helena A Woroniecka, Richard Brust, Aya Sabbah, Michael Towrie, et al. Enhancing proton-coupled electron transfer in blue light using fad photoreceptor appabluf. Journal of the American Chemical Society, 147(1):39–44, 2024.

[31] Berk Hess, Henk Bekker, Herman JC Berendsen, and Johannes GEM Fraaije. LINCS: a linear constraint solver for molecular simulations. J. Comput. Chem., 18(12):1463–1472, 1997.

[32] LiHong Hu, Pär Söderhjelm, and Ulf Ryde. On the convergence of QM/MM energies. J. Chem. Theor. Comput., 7(3):761–777, 2011.

[33] Jing Huang and Alexander D MacKerell Jr. CHARMM36 all-atom additive protein force field: Validation based on comparison to NMR data. J. Comput. Chem., 34(25):2135–2145, 2013.

[34] Jürg Hutter, Marcella Iannuzzi, Florian Schiffmann, and Joost VandeVondele. cp2k: atomistic simulations of condensed matter systems. Wiley Interdiscip. Rev. Comput. Mol. Sci., 4(1):15–25, 2014.

[35] Mineo Iseki, Shigeru Matsunaga, Akio Murakami, Kaoru Ohno, Kiyoshi Shiga, Kazuichi Yoshida, Michizo Sugai, Tetsuo Takahashi, Terumitsu Hori, and Masakatsu Watanabe. A blue-light-activated adenylyl cyclase mediates photoavoidance in euglena gracilis. Nature, 415(6875):1047–1051, 2002.

[36] Tatsuya Iwata, Takashi Nagai, Shota Ito, Shinsuke Osoegawa, Mineo Iseki, Masakatsu Watanabe, Masashi Unno, Shinya Kitagawa, and Hideki Kandori. Hydrogen Bonding Environments in the Photocycle Process around the Flavin Chromophore of the AppA-BLUF domain. J. Am. Chem. Soc., 140(38):11982–11991, 2018.

[37] Christopher Jarzynski. Nonequilibrium equality for free energy differences. Phys. Rev. Lett, 78(14):2690, 1997.

[38] W. L. Jorgensen, J. Chandrasekhar, J. D. Madura, R. W. Impey, and M. L. Klein. Comparison of simple potential functions for simulating liquid water. J. Chem. Phys., 79:926–935, 1983.

[39] Astrid Jung, Tatiana Domratcheva, Marina Tarutina, Qiong Wu, Wen-huang Ko, Robert L Shoeman, Mark Gomelsky, Kevin H Gardner, and Ilme Schlichting. Structure of a bacterial BLUF photoreceptor: insights into blue light-mediated signal transduction. Proc. Natl. Acad. Sci. USA, 102(35):12350–12355, 2005.

[40] Astrid Jung, Jochen Reinstein, Tatiana Domratcheva, Robert L Shoeman, and Ilme Schlichting. Crystal structures of the AppA BLUF domain photoreceptor provide insights into blue light-mediated signal transduction. J. Mol. Biol., 362(4):717–732, 2006.

[41] Kristof Karadi, Sofia M Kapetanaki, Katalin Raics, Ildiko Pecsi, Robert Kapronczai, Zsuzsanna Fekete, James N Iuliano, Jinnette Tolentino Collado, Agnieszka A Gil, Jozsef Orban, et al. Functional dynamics of a single tryptophan residue in a BLUF protein revealed by fluorescence spectroscopy. Sci. Rep., 10(1):1–15, 2020.

[42] Maria G Khrenova, Tatiana Domratcheva, and Alexander V Nemukhin. Molecular mechanism of the dark-state recovery in BLUF photoreceptors. Chem. Phys. Lett., 676:25–31, 2017.

[43] Maria G Khrenova, Alexander V Nemukhin, and Tatiana Domratcheva. Photoinduced electron transfer facilitates tautomerization of the conserved signaling glutamine side chain in BLUF protein light sensors. J. Phys. Chem. B, 117(8):2369–2377, 2013.

[44] Murat Kılıç and Bernd Ensing. Redox properties of flavin in bluf and lov photoreceptor proteins from hybrid qm/mm molecular dynamics simulation. The Journal of Physical Chemistry B, 128(13):3069–3080, 2024.

[45] Akiko Kita, Koji Okajima, Yukio Morimoto, Masahiko Ikeuchi, and Kunio Miki. Structure of a cyanobacterial BLUF protein, Tll0078, containing a novel FAD-binding blue light sensor domain. J. Mol. Biol., 349(1):1–9, 2005.

[46] Brian J Kraft, Shinji Masuda, Jun Kikuchi, Vladimira Dragnea, Gordon Tollin, Jeffrey M Zaleski, and Carl E Bauer. Spectroscopic and mutational analysis of the blue-light photoreceptor AppA: a novel photocycle involving flavin stacking with an aromatic amino acid. Biochemistry, 42(22):6726–6734, 2003.

[47] Thomas D Kühne, Marcella Iannuzzi, Mauro Del Ben, Vladimir V Rybkin, Patrick Seewald, Frederick Stein, Teodoro Laino, Rustam Z Khaliullin, Ole Schütt, Florian Schiffmann, et al. CP2K: An electronic structure and molecular dynamics software package-Quickstep: Efficient and accurate electronic structure calculations. J. Chem. Phys., 152(19):194103, 2020.

[48] Wouter Laan, Michael A van der Horst, Ivo H Van Stokkum, and Klaas J Hellingwerf. Initial Characterization of the Primary Photochemistry of AppA, a Blue-light–using Flavin Adenine Dinucleotide–domain Containing Transcriptional Antirepressor Protein from Rhodobacter sphaeroides: A Key Role for Reversible Intramolecular Proton Transfer from the Flavin Adenine Dinucleotide Chromophore to a Conserved Tyrosine? Photochem. Photobiol., 78(3):290–297, 2003.

[49] Andras Lukacs, Peter J Tonge, and Stephen R Meech. Photophysics of the blue light using flavin domain. Acc. Chem. Res., 55(3):402–414, 2022.

[50] Shinji Masuda and Carl E Bauer. AppA is a blue light photoreceptor that antirepresses photosynthesis gene expression in Rhodobacter sphaeroides. Cell, 110(5):613–623, 2002.

[51] Shinji Masuda, Koji Hasegawa, Hiroyuki Ohta, and Taka-aki Ono. Crucial role in light signal transduction for the conserved Met93 of the BLUF protein PixD/Slr1694. Plant Cell Physiol., 49(10):1600–1606, 2008.

[52] Shinji Masuda, Koji Hasegawa, and Taka-aki Ono. Tryptophan at position 104 is involved in transforming light signal into changes of β -sheet structure for the signaling state in the BLUF domain of AppA. Plant Cell Physiol., 46(12):1894–1901, 2005.

[53] Shinji Masuda, Yoshiyuki Tomida, Hiroyuki Ohta, and Ken-ichiro Takamiya. The critical role of a hydrogen bond between Gln63 and Trp104 in the blue-light sensing BLUF domain that controls AppA activity. J. Mol. Biol., 368(5):1223–1230, 2007.

[54] Tilo Mathes, Ivo HM van Stokkum, Manuela Stierl, and John TM Kennis. Redox modulation of flavin and tyrosine determines photoinduced proton-coupled electron transfer and photoactivation of BLUF photoreceptors. J. Biol. Chem., 287(38):31725–31738, 2012.

[55] Katharina Meier, Walter Thiel, and Wilfred F van Gunsteren. On the effect of a variation of the force field, spatial boundary condition and size of the QM region in QM/MM MD simulations. J. Comput. Chem., 33(4):363–378, 2012.

[56] Maria-Andrea Mroginski, Suliman Adam, Gil S Amoyal, Avishai Barnoy, Ana-Nicoleta Bondar, Veniamin Borin, Jonathan R Church, Tatiana Domratcheva, Bernd Ensing, Francesca Fanelli, et al. Frontiers in multiscale modelling of photoreceptor proteins. Photochem. Photobiol., 97(2):243–269, 2021.

[57] M. Parrinello and A. Rahman. Polymorphic transitions in single crystals: A new molecular dynamics method. J. Appl. Phys., 52:7182–7190, 1981.

[58] John P Perdew, Kieron Burke, and Matthias Ernzerhof. Generalized gradient approximation made simple. Phys. Rev. Lett., 77(18):3865, 1996.

[59] A Pèrez de Alba Ortíz, A Tiwari, RC Puthenkalathil, and B Ensing. Advances in enhanced sampling along adaptive paths of collective variables. J. Chem. Phys., 149(7):072320, 2018.

[60] Alberto Pèrez de Alba Ortíz, Jocelyne Vreede, and Bernd Ensing. The adaptive path collective variable: a versatile biasing approach to compute the average transition path and free energy of molecular transitions. In Massimiliano Bonomi and Carlo Camilloni, editors, Biomolecular Simulation, pages 255–290. Springer, 2019.

[61] Keyarash Sadeghian, Marco Bocola, and Martin Schütz. A conclusive mechanism of the photoinduced reaction cascade in blue light using flavin photoreceptors. J. Am. Chem. Soc., 130(37):12501–12513, 2008.

[62] Giacomo Salvadori, Patrizia Mazzeo, Davide Accomasso, Lorenzo Cupellini, and Benedetta Mennucci. Deciphering photoreceptors through atomistic modeling from light absorption to conformational response. Journal of molecular biology, 436(5):168358, 2024.

[63] Elvira R Sayfutyarova, Joshua J Goings, and Sharon Hammes-Schiffer. Electron-Coupled Double Proton Transfer in the Slr1694 BLUF Photoreceptor: A Multireference Electronic Structure Study. J. Phys. Chem. B, 123(2):439–447, 2018.

[64] Hans Martin Senn and Walter Thiel. QM/MM methods for biomolecular systems. Angew. Chem. Int. Edit., 48(7):1198–1229, 2009.

[65] Masahiko Taguchi, Shun Sakuraba, Justin Chan, and Hidetoshi Kono. Unveiling the photoactivation mechanism of bluf photoreceptor protein through hybrid quantum mechanics/molecular mechanics free-energy calculation. ACS Physical Chemistry Au, 4(6):647–659, 2024.

[66] Shunrou Tokonami, Morihiko Onose, Yusuke Nakasone, and Masahide Terazima. Slow conformational changes of blue light sensor bluf proteins in milliseconds. J. Am. Chem. Soc., 144(9):4080–4090, 2022.

[67] Jing Tong, Peng Zhang, Lei Zhang, Dongwei Zhang, David N Beratan, Haifeng Song, Yi Wang, and Tie Li. A robust bioderived wavelength-specific photosensor based on BLUF proteins. Sens. Actuators B Chem., 310:127838, 2020.

[68] Anikó Udvarhelyi and Tatiana Domratcheva. Photoreaction in BLUF Receptors: Proton-coupled Electron Transfer in the Flavin-Gln-Tyr System. Photochem. Photobiol., 87(3):554–563, 2011.

[69] Anikó Udvarhelyi and Tatiana Domratcheva. Glutamine rotamers in BLUF photoreceptors: a mechanistic reappraisal. J. Phys. Chem. B, 117(10):2888–2897, 2013.

[70] Anikó Udvarhelyi, Massimo Olivucci, and Tatiana Domratcheva. Role of the molecular environment in flavoprotein color and redox tuning: QM cluster versus QM/MM modeling. J. Chem. Theor. Comput., 11(8):3878–3894, 2015.

[71] Michael A Van Der Horst and Klaas J Hellingwerf. Photoreceptor proteins, “star actors of modern times”: a review of the functional dynamics in the structure of representative members of six different photoreceptor families. Acc. Chem. Res., 37(1):13–20, 2004.

[72] David Van Der Spoel, Erik Lindahl, Berk Hess, Gerrit Groenhof, Alan E Mark, and Herman JC Berendsen. GROMACS: fast, flexible, and free. J. Comput. Chem., 26(16):1701–1718, 2005.

[73] Arieh Warshel and Michael Levitt. Theoretical studies of enzymic reactions: dielectric, electrostatic and steric stabilization of the carbonium ion in the reaction of lysozyme. J. Mol. Biol., 103(2):227–249, 1976.

[74] Yang Xu, Peng Bao, Kai Song, and Qiang Shi. Theoretical study of proton coupled electron transfer reaction in the light state of the AppA BLUF photoreceptor. J. Comput. Chem., 40(9):1005–1014, 2019.

[75] Yalin Zhou, Xiu-Wen Kang, Zhongneng Zhou, Zijing Chen, Shuhua Zou, Siwei Tang, Bingyao Wang, Kailin Wang, Dongping Zhong, and Bei Ding. Unified mechanism of light-state bluf domain photocycles by capturing proton relay intermediates. Ultrafast Science, 4:0072, 2024.

[76] Peyman Zirak, Alfons Penzkofer, T Schiereis, Peter Hegemann, Astrid Jung, and Ilme Schlichting. Photodynamics of the small BLUF protein BlrB from Rhodobacter sphaeroides. Photochem. Photobiol. B, Biol., 83(3):180–194, 2006.

